# Inhibiting Host Protein Deposition on Urinary Catheters Reduces Urinary Tract Infections

**DOI:** 10.1101/2021.12.07.471578

**Authors:** Marissa Jeme Andersen, ChunKi Fong, Alyssa Ann La Bella, Alex Molesan, Matthew M. Champion, Caitlin Howell, Ana L. Flores-Mireles

## Abstract

Microbial adhesion to medical devices is common for hospital-acquired infections, particularly for urinary catheters. If not properly treated these infections cause complications and exacerbate antimicrobial resistance. Catheter use elicits bladder inflammation, releasing host serum-proteins, including fibrinogen, into the bladder, which deposit on the urinary catheter. *Enterococcus faecalis* uses fibrinogen as a scaffold to bind and persist in the bladder despite antibiotic treatments. Inhibition of fibrinogen-pathogen interaction significantly reduces infection. Here, we show deposited fibrinogen is advantageous for uropathogens, suggesting that targeting catheter protein deposition may reduce colonization creating an effective intervention. Host-protein deposition was reduced, using liquid-infused catheters, resulting in decreased colonization on catheters, in bladders, and dissemination *in vivo*. Furthermore, proteomics revealed a significant decrease in deposition of host-secreted proteins on liquid-infused catheter surfaces. Our findings suggest targeting microbial binding scaffolds may be an effective antibiotic-sparing intervention for use against catheter-associated urinary tract infections and other medical device infections.

## Main Text

Urinary catheters drain patient’s bladders during surgical sedation and recovery in addition to being used in treatment for a variety of conditions, making it an exceedingly common procedure in healthcare facilities. Despite the benefits urinary catheters provide for patients, catheterization causes adverse effects including infections and bladder stones (Andersen & Flores-Mireles, 2019; Feneley, Hopley, & Wells, 2015). The most common complication is catheter-associated urinary tract infection (CAUTI), which accounts for 40% of all hospital acquired infections (HAIs) (Andersen & Flores-Mireles, 2019; Feneley et al., 2015). Unfortunately, CAUTIs often lead to bloodstream infections and systemic dissemination with a 30% mortality rate, causing significant financial burdens for hospitals and patients (Andersen & Flores-Mireles, 2019; Feneley et al., 2015).

New and current management guidelines for CAUTIs have resulted in moderate reductions in incidences (Assadi, 2018; Meddings et al., 2014). The standard treatment for patients with symptomatic CAUTI is catheter removal and replacement and an antibiotic regiment (Assadi, 2018; A. Flores-Mireles, Hreha, & Hunstad, 2019). However, this approach isn’t effective because biofilms on the catheter surface protect microbes against antibiotics and the immune system and the potential development of antimicrobial resistance (A. Flores-Mireles et al., 2019; Trautner & Darouiche, 2004). Many ongoing efforts to prevent biofilms on catheters are focused on developing surface modifications (Andersen & Flores-Mireles, 2019; Faustino, Lemos, Monge, & Ribeiro, 2020; Singha, Locklin, & Handa, 2017). Catheters impregnated with antimicrobials, such as metal ions and antibiotics, have become popular and are now commercialized due to promising *in vitro* work but, in clinical trials these catheters have shown, at best, mixed results (Andersen & Flores-Mireles, 2019; Singha et al., 2017). Importantly, there is concern that this approach may not be a long-term solution given that the presence of antimicrobial compounds may drive development of resistance (Singha et al., 2017; Westfall et al., 2019).

Recent findings have unveiled the importance of the host clotting factor 1, fibrinogen (Fg), for surface adhesion and subsequent establishment of biofilms and persistence of CAUTIs in *Enterococcus faecalis* and *Staphylococcus aureus* infections (A. Flores-Mireles et al., 2019; Gaston et al., 2020; Klein & Hultgren, 2020). Fg is continuously released into the bladder lumen in response to mechanical damage to the urothelial lining caused by catheterization (A. Flores-Mireles et al., 2019; Klein & Hultgren, 2020). Once in the lumen, Fg is deposited on the catheter, providing a scaffold for these incoming uropathogens to bind to and when blocking the interaction between Fg and *E. faecalis* using antibodies, the pathogen is not able to effectively colonize the bladder (Di Venanzio et al., 2019; A. L. Flores-Mireles, Pinkner, Caparon, & Hultgren, 2014; A. L. Flores-Mireles et al., 2016; Gaston et al., 2020; Walker et al., 2017).

Thus, we hypothesized that reducing availability of binding scaffolds, in this case Fg, would decrease microbial colonization in a catheterized bladder. To test our hypothesis, we used a mouse model of CAUTI and a diverse panel of uropathogens, including *E. faecalis, C. albicans*, uropathogenic *Escherichia coli, Pseudomonas aeruginosa, A. baumannii*, and *Klebsiella pneumonia*, which we found all bind more extensively to catheters with Fg present. To resolve the deposition of Fg, we focus on anti-fouling modifications, specifically, liquid infused silicone (LIS). LIS is more simple to make, more stable and more cost effective than other anti-fouling polymer modifications (Andersen & Flores-Mireles, 2019; Campoccia, Montanaro, & Arciola, 2013; Homeyer, Goudie, Singha, & Handa, 2019; Howell, Grinthal, Sunny, Aizenberg, & Aizenberg, 2018; Sedlarik, 2013; Singha et al., 2017; Sotiri et al., 2018; Villegas, Zhang, Abu Jarad, Soleymani, & Didar, 2019). Additionally, LIS has shown to reduce clotting in central lines and infection in skin implants (Chen et al., 2017; Leslie et al., 2014). We show that our LIS-catheters reduced Fg deposition and microbial binding not only *in vitro* but also *in vivo*. Furthermore, LIS-catheters significantly decrease host-protein deposition when compared to unmodified (UM)-catheters as well as reducing catheter-induced inflammation. These findings suggest that targeting host-protein deposition on catheter surfaces and the use of LIS-catheters are plausible strategies for reducing instances of CAUTI.

## Results

### Uropathogens interact with Fg during CAUTI

Due to the understood interaction between Fg and some uropathogens as well as Fg accumulation on catheters over time in human and mice, we assessed potential interaction of *E. faecalis* OG1RF (positive control) uropathogenic *E. coli* UTI89, *P. aeruginosa* PAO1, *K. pneumoniae* TOP52, *A. baumannii* UPAB1, and *C. albicans* SC5314 with Fg *in vivo*, using a CAUTI mouse model (A. Flores-Mireles et al., 2019; A. L. Flores-Mireles et al., 2014; A. L. Flores-Mireles et al., 2016). Mice catheterized and infected with the respective uropathogen were sacrificed at 24 post infection (hpi). Catheters and bladders were harvested, stained and imaged. Visual and quantitative analysis of the catheters showed all uropathogens co-localizing strongly with Fg deposits exhibiting preference for Fg (**Figure 1A and B**) and robust Fg deposition on catheters, validating previous studies on human catheters (A. Flores-Mireles et al., 2019; A. L. Flores-Mireles et al., 2014; A. L. Flores-Mireles et al., 2016). Importantly, immunofluorescence (IF) analysis of bladder sections showed that all uropathogens interact with Fg on the bladder urothelium or in the lumen during CAUTI (**Figure 1C** and montages in **Figure S1**). Although we show interaction between the pathogens and Fg, further studies are needed to characterize each pathogen-Fg interaction mechanism, as previously done with *E. faecalis* and *S. aureus* (A. L. Flores-Mireles et al., 2014; A. L. Flores-Mireles et al., 2016; Walker et al., 2017).

**Figure 1.**
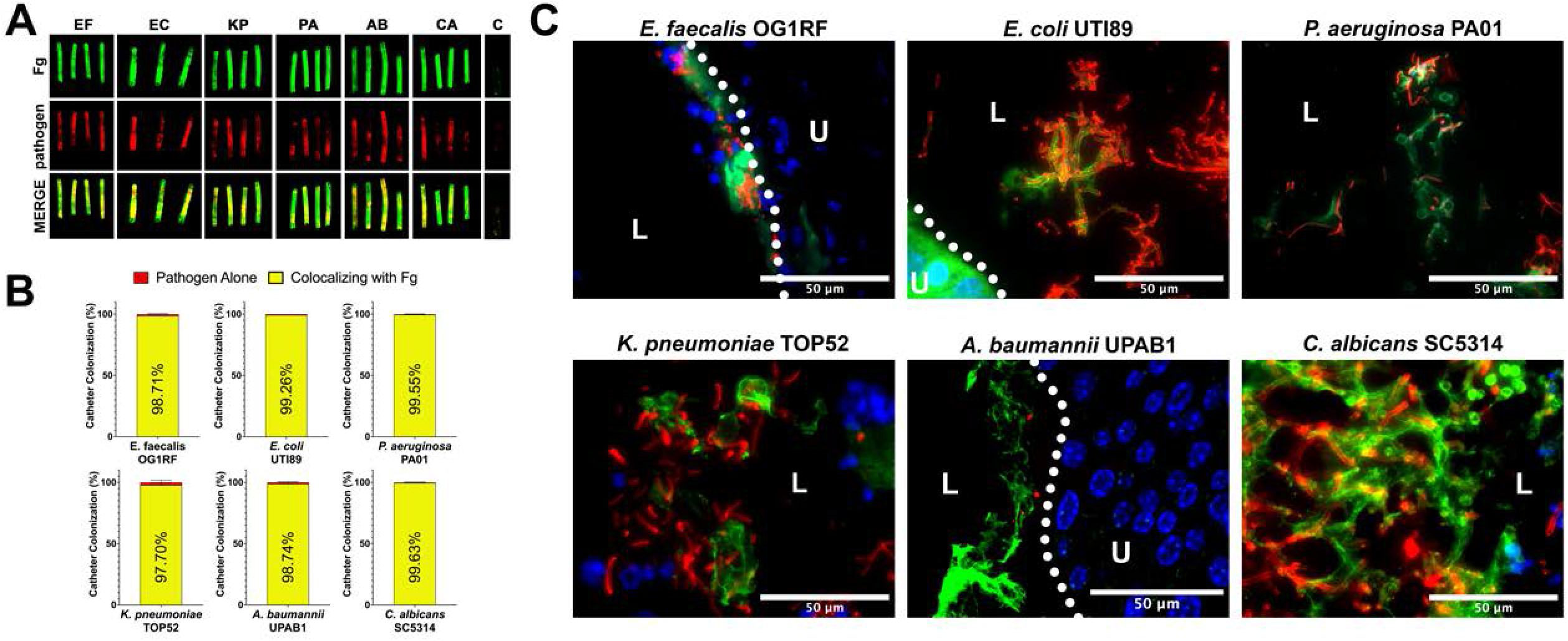
Uropathogens interact with Fg *in vivo*. **A)** Urinary catheters stained with IF for Fg deposition (Fg; green) and microbe binding (respective pathogen; red). Un-implanted catheters were used as controls for autofluorescence, n=3-4. **B)** Quantification of uropathogen-Fg colocalization on catheters from panel a. **C)** Representative images from a single bladder illustrating the interaction of uropathogens (red), Fg (green) and nuclei (blue) on the bladder urothelium (U) and in the lumen (L). Scale bars = 50 nm. Montages can be found in supplemental figure 1. For all graphs error bars show the standard error of the mean (SEM). Between 3-5 replicates of n=4-12 each were performed for each pathogen and condition.

### Fg on urinary catheter material enhances microbial binding

Based on our *in vivo* findings, we assessed whether Fg could promote initial binding of the uropathogens to silicone catheters as previously seen for *E. faecalis* (A. L. Flores-Mireles et al., 2014). In addition to Fg, bovine serum albumin (BSA) was tested since serum albumin is one of the most abundant protein on human and mouse urinary catheters (Molina, In preparation) (**Table S1**). We compared uropathogen binding to Fg-, BSA- and uncoated silicone, finding that Fg significantly enhanced the binding to the catheter for all uropathogens when compared with uncoated and BSA-coated silicone catheters (**Figure 2**). Interestingly, *P. aeruginosa* and *A. baumannii* binding to BSA-coated silicone was ~14% and ~10% higher than uncoated controls, respectively (**Figure 2C and E**), illuding to a role for other host-secreted proteins during infection. However, these values were still significantly lower than the increase in binding observed on Fg-coated silicone (**Figure 2C and E**). Taken together, these data suggest that uropathogen interaction with host-proteins deposited on silicone surfaces, particularly Fg, increases the ability of uropathogens to colonize urinary catheters.

**Figure 2.**
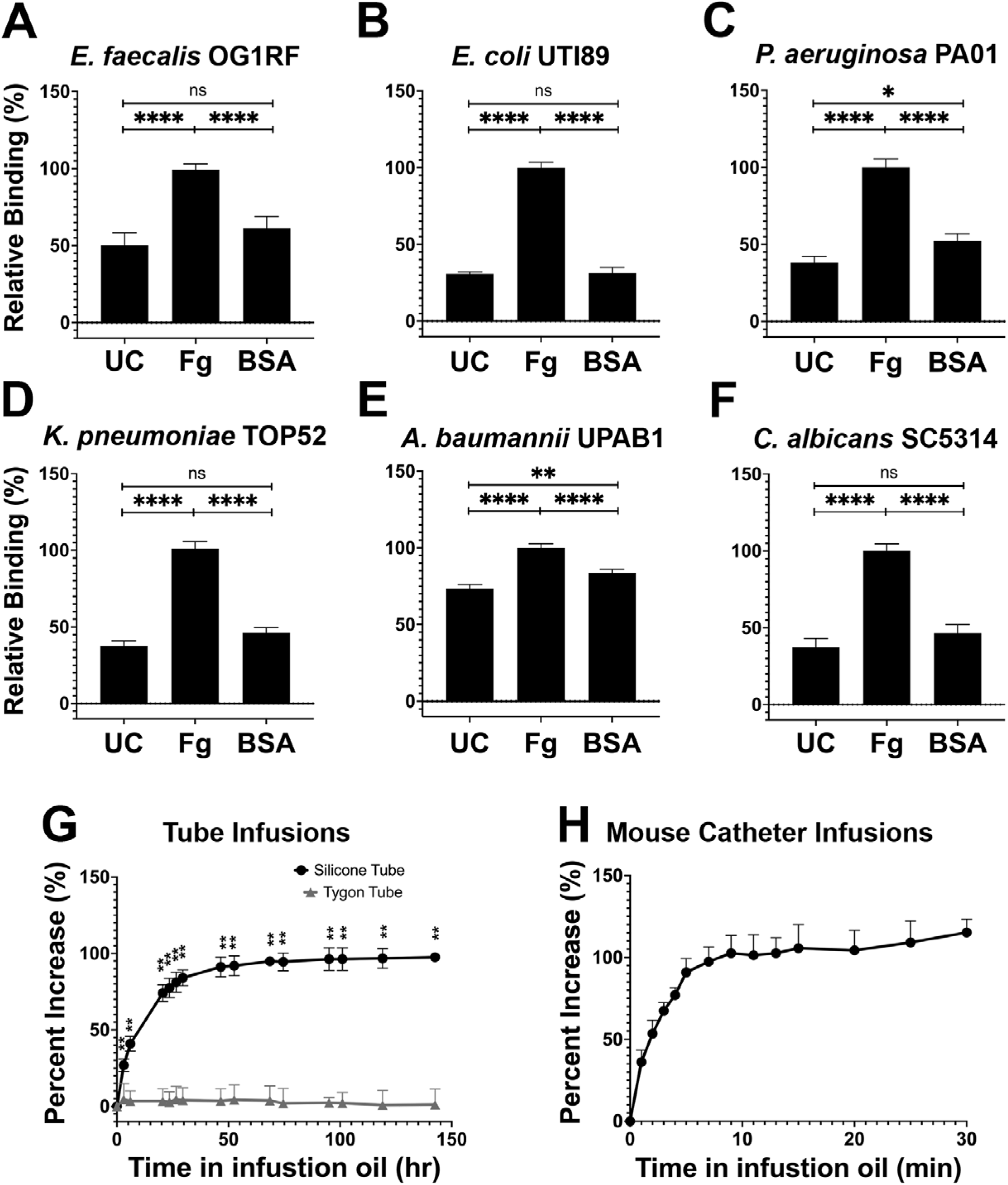
Silicone infusion and Fg enhancement of microbial surface binding. **A-F)** Uropathogens were tested for their ability to bind to protein coated and uncoated (UC) silicone catheters. For all graphs error bars show the standard error of the mean (SEM). Between 3-5 replicates of n=4-12 each were performed for each pathogen and condition. **G)** Kinetics of silicone oil infusion on silicone and Tygon tubes, as well as **H)** mouse silicone catheters.

### Characterization of liquid-infused catheters to prevent host-protein deposition

Based on the exploitative interaction of uropathogens with deposited Fg, we hypothesized that development of a material to prevent protein deposition would also reduce microbial colonization. Recent work with liquid-infused surfaces have demonstrated resistance to protein and bacterial fouling (Goudie, Pant, & Handa, 2017; Howell et al., 2018; Leslie et al., 2014; Sotiri et al., 2018). This prompted us to develop a LIS material by modifying medical-grade silicone using inert trimethyl-terminated polydimethylsiloxane fluid (silicone oil) (Goudie et al., 2017). Analysis of the oil’s infusion rate showed a significant increase in silicone weight during the first 3 days of infusion then a gradual decrease in infusion until a plateau was reached after ~50 hrs (raw weight in **Figure S3** and; **Figure S2D** in black). Plastic Tygon tubes (non-silicone) were used as negative controls (**Figure S2D** in grey). Full infusion of mouse silicone catheters was achieved by 10 min (**Figure S2C and E**). Investigation of silicone tube dimensions showed an increase in length, outer and inner diameter of ~41.3%, ~103.1%, and ~27.6%, respectively (**Figure S2F**) and mouse catheters showed an increase of ~30.7%, ~28.7%, and ~39.8%, respectively (**Figure S2G**).

### LIS modification reduces Fg deposition and microbial binding in vitro

The ability of the LIS-catheters to reduce Fg deposition *in vitro* was tested for infused medical-grade silicone material and two commercially available urinary catheters, Dover and Bardex with unmodified (UM) versions of each used as controls. Each was incubated with Fg overnight and assessed by IF. We found that Fg deposition was reduced in all LIS-catheters, showing ~90% decrease on Dover and ~100% on the Bardex and medical-grade silicone tubing when compared with the corresponding UM controls (**Figure 3A,B**).

**Figure 3.**
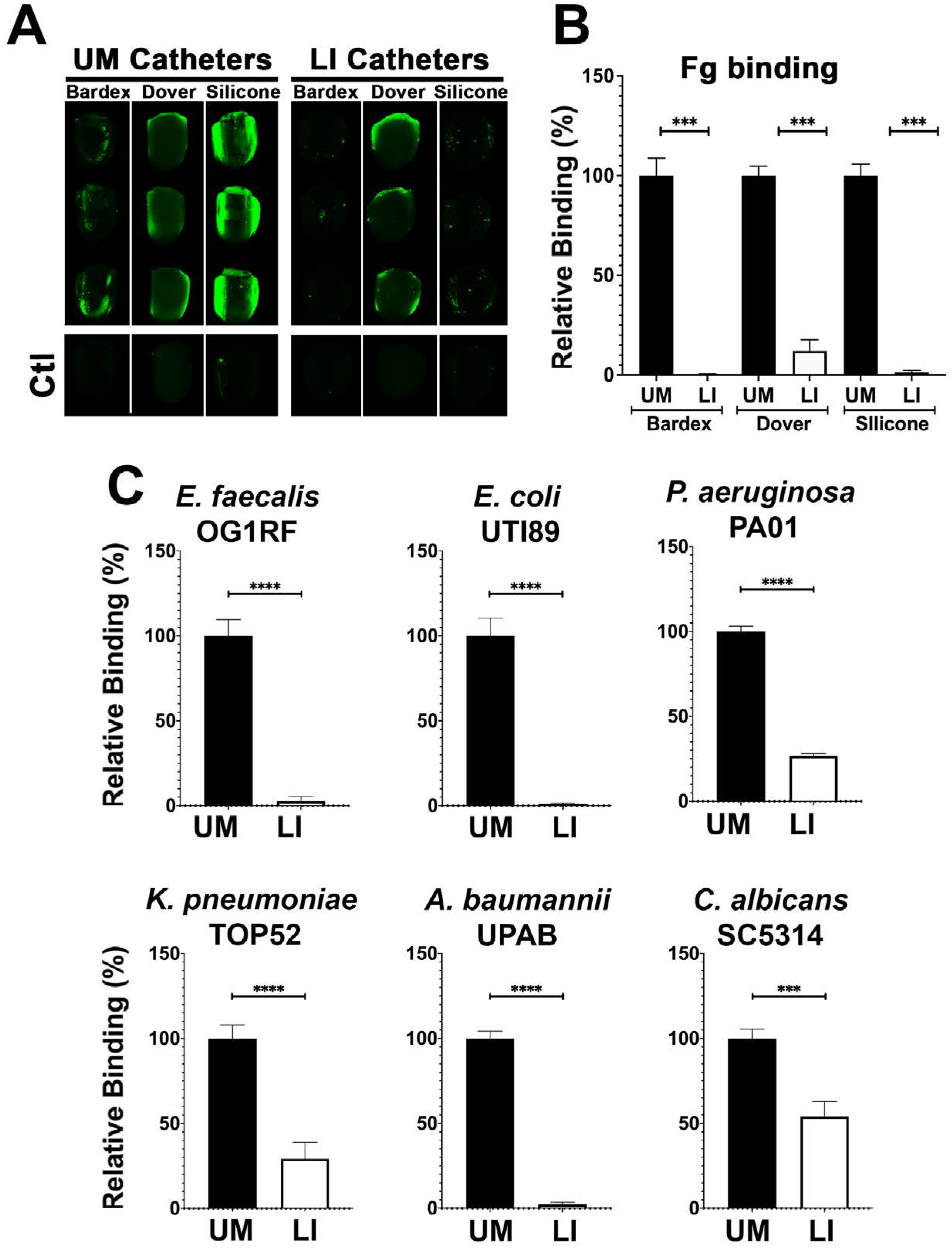
LIS-modification reduces Fg deposition and microbial binding *in vitro*. **A)** Visualization and **B)** quantification of Fg (green) deposition on UM-catheter material (black bars) and LIS-catheter materials (white bars) by IF staining. 3 replicates with n=2-3 each. **C)** Microbial binding on unmodified (UM) and LIS. For all graphs error bars show the standard error of the mean (SEM) and each condition had 3 replicates with n=3 each.

Based on previous reports of the biofouling ability of liquid-infused surfaces and our LIS’s success in reducing Fg deposition, we tested its ability to prevent microbial surface binding (Howell et al., 2018). Our six uropathogens were grown in urine supplemented with BSA at 37°C (**Table S2**), cultures were normalized in urine, added to UM control and LIS-catheters, incubated under static conditions and assessed via IF. Analysis found the LIS-catheters showed significantly reduced binding of all uropathogens when compared to UM controls (**Figure 3C**). These results further demonstrate the capability of silicone LIS-catheters to reduce not only protein deposition but to also impede microbial colonization.

### Fg deposition and microbial biofilms on catheters was reduced by LIS

Mice were catheterized with either an UM-or LIS-catheter and infected with one of six uropathogens for 24hrs. Bladders and catheters were harvested and assessed for microbial burden by CFU enumeration or fixed for staining. Kidneys, spleens and hearts were collected to determine microbial burden. We found that mice with LIS-catheters significantly reduced microbial colonization in the bladder and on catheters when compared with UM-catheterized mice regardless of the infecting uropathogen (**Figure 4**). Additionally, colonization was significantly lower in LIS-catheterized mouse kidneys for *P. aeruginosa A. baumannii*, and *E. coli* infections (**Figure 4C, E, and G**) and LIS-catheterized mice infected with *E. coli* or *C. albicans* showed significantly less colonization of the spleen (**Figure 4G and K**). *K. pneumoniae* kidney and spleen colonization was not statistically significant, however they showed a trend of less colonization and significantly less colonization of the heart **(Figure 4I)**.

**Figure 4.**
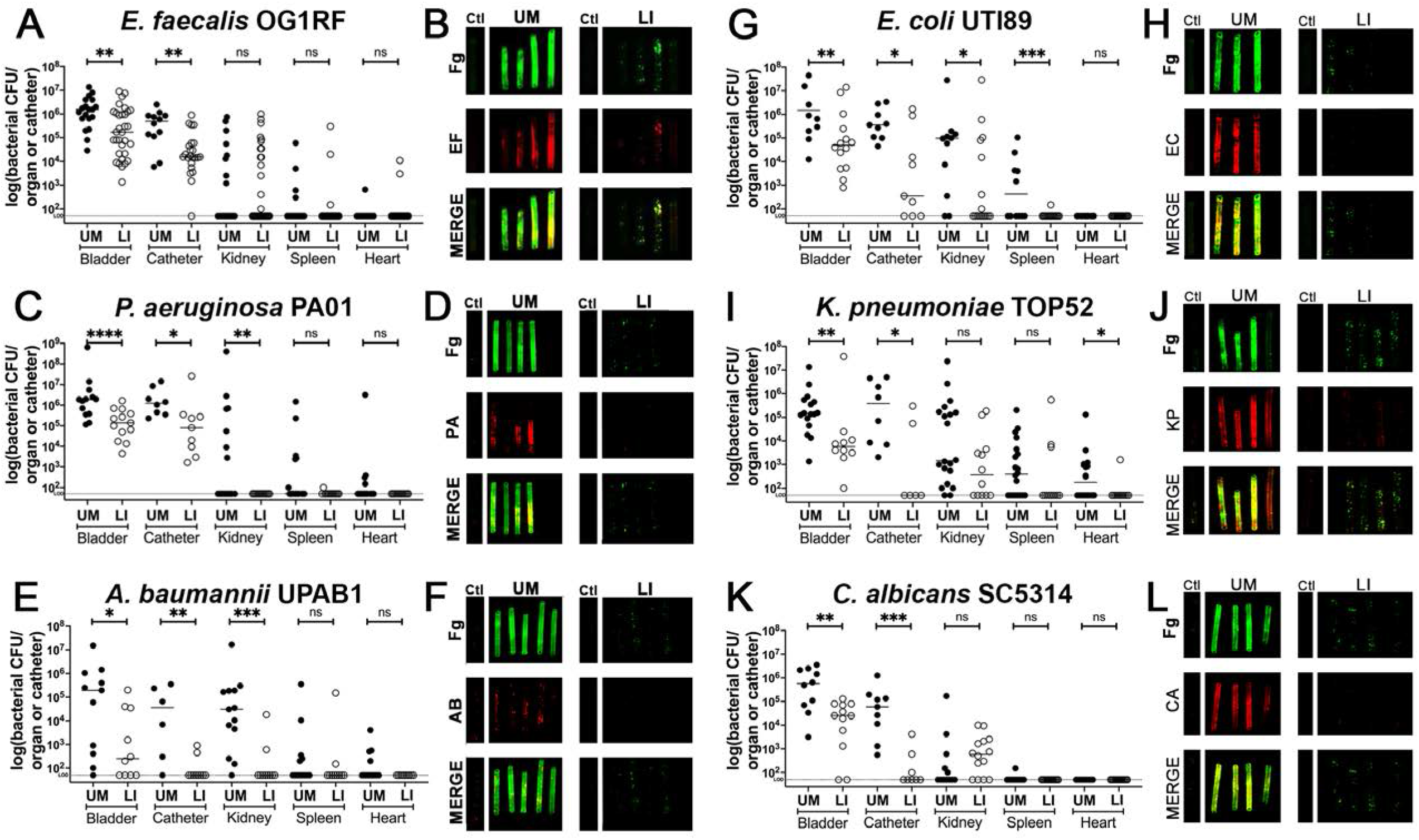
*In vivo* reduction of Fg deposition to restrict microbial burden. Mice were catheterized and infected with one of six uropathogens. **A,C,E,G,I,K)** Organ and catheter CFUs from mice with either an UM-catheter (closed circles) or LIS-catheter (open circles) show the dissemination profile of the pathogen. **B,D,F,G,I,K)** Imaging of catheters for Fg (green), respective uropathogen (red), and a merged image compare deposition on UM-catheters (left) with LIS-catheters (right); non-implanted catheters as controls. Quantification of microbial colonization and co-localization on the catheters can be found in supplemental figure 2. All animal studies for CFUs, catheter and bladder imaging had at least 10 animals per strain and catheter type.

Furthermore, IF imaging and quantification of catheters confirmed decreased Fg deposition and microbial biofilms on LIS-catheters compared to UM (**Figure 4B, D, F, H, J, I** and **S3**). These data demonstrate pathogens preferentially bind to Fg, and that the LIS-modification successfully reduced Fg deposition (the microbes’ binding platform), disrupts uropathogen biofilm formation on catheters, and colonization of the bladder *in vivo*. Importantly, H&E analysis shows the LI-catheter does not exacerbate bladder inflammation regardless of the presence of infection or not (**Figure 5A-G**), an important factor to account for when developing a new medical device. In fact, for some pathogens, the LI-catheter results in less inflammation than bladders catheterized with an UM-catheter (**Figure 5B, C, and G**). Furthermore, we examine Fg presence, uropathogen colonization, and neutrophil recruitment in UM- and LIS-catheterized and infected bladders by IF microscopy. This analysis revealed a reduction of microbial colonization as well as decreased neutrophil recruitment (**Figure 5H-M**).

**Figure 5.**
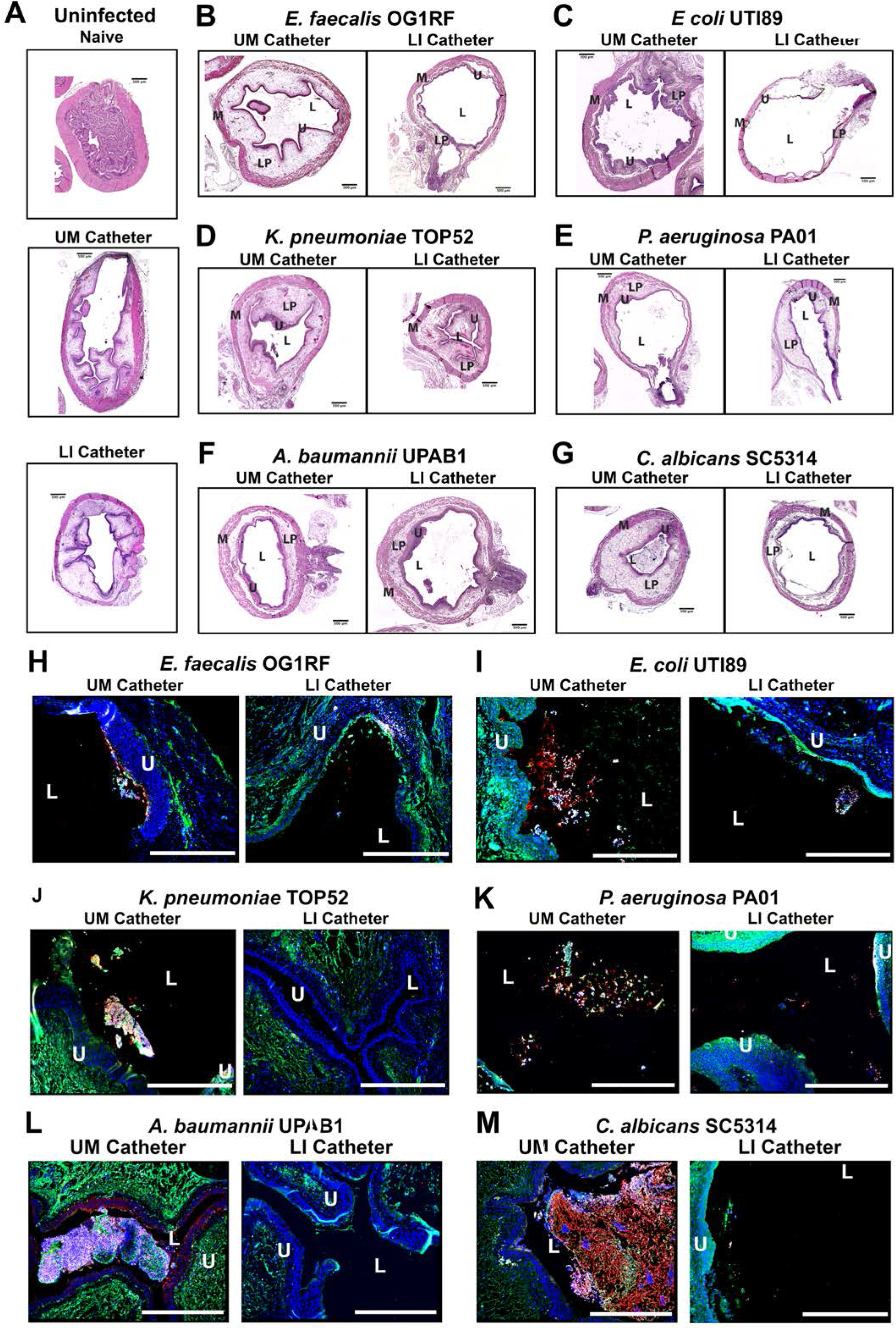
LIS-catheters reduce bladder colonization and inflammation. Mice were catheterized and inoculated one of six strains. **A)** Naïve or bladders catheterized with an UM or LIS-catheter were uninfected controls. **B-G)** Bladder sections were stained with H&E to compare inflammation from UM-catheters (left) and LIS-catheters (right). **C-M)** 20x images of IF stained bladders catheterized with an UM-catheter (left panels) or a LIS-catheter (right panels). Bladders stained for nuclei (blue), Fg (green), respective uropathogens (red) and neutrophils (white). The urothelial/lumen boundaries are outlined in white dotted lines and labeled U (urothelium) and L (lumen) and all scale bars are 500um. Montages can be found in supplemental figure 3.

### LIS modification reduces protein deposition on catheters in CAUTI mouse model of E. faecalis

A quantitative-proteomics comparison was performed to identify proteins deposited on UM- and LIS-catheters retrieved 24hpi with *E. faecalis*. Harvested catheters were prepared and protease digested with trypsin as in Zougman et al. (2014) nLC-MS/MS was performed in technical duplicate and label-free-proteomics (LFQ) processed as in Cox and Mann, 865 proteins were identified at a 1% FDR (Cox & Mann, 2008). Total abundance of protein was significantly reduced in LIS-catheters *vs* UM-catheters (**Figure 6A**). Additionally, abundance of Fg and over 130 other proteins significantly decreased while only three proteins showed a significant increase (UDP-glucose 6-dehydrogenase, filamin-B, and proteasome subunit beta type-5) (**Figure 6B**). This data further demonstrates that the LIS modification not only reduced Fg deposition but also a wide variety of host-proteins, which could play a role in microbial colonization and biofilm formation as demonstrated earlier with BSA (**Figure 2C and E**)

**Figure 6.**
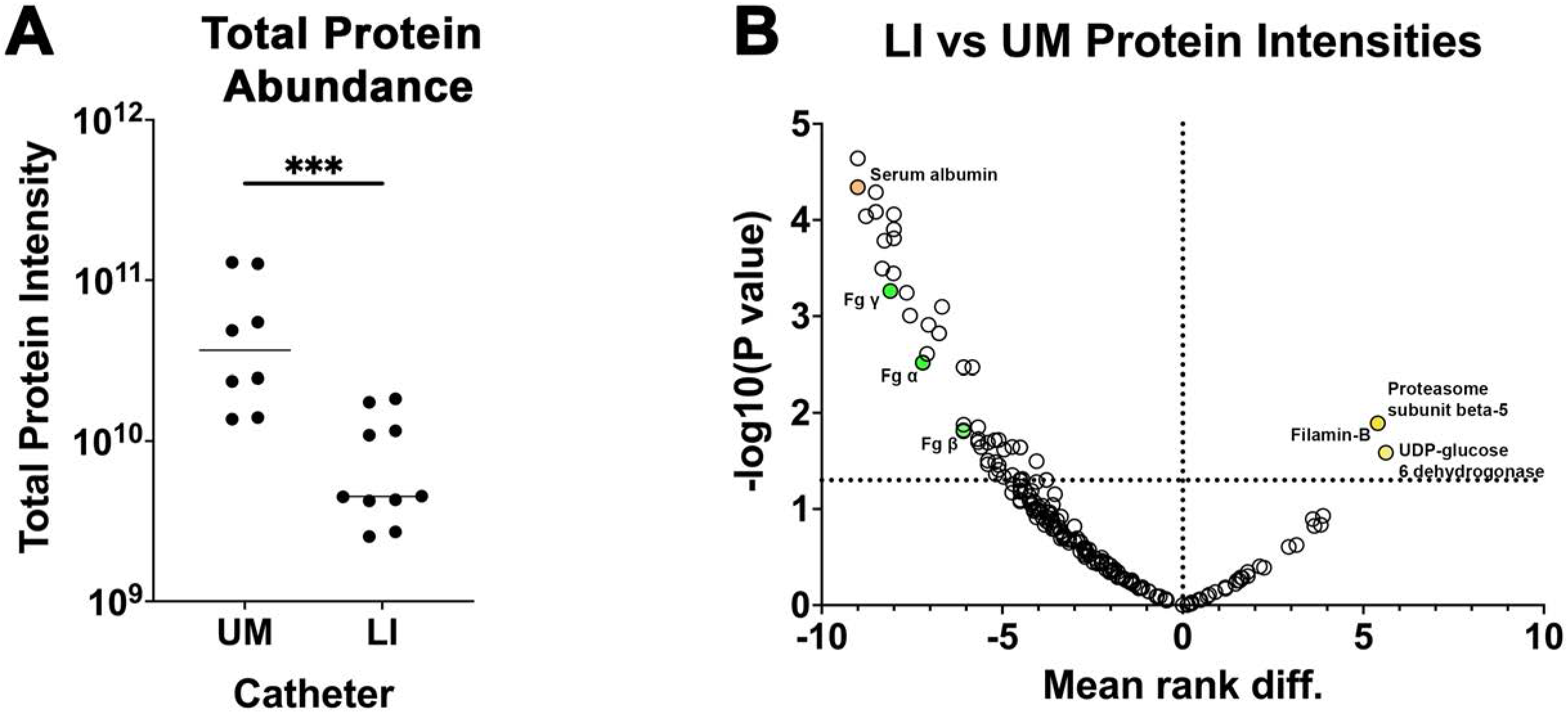
LIS-catheter reduces host-protein deposition *in vivo*. A subset of UM-catheters and LIS-catheters taken from mice 24 hpi with *E. faecalis* were assessed for protein deposition via mass spectrometry 4-UM catheters and 5 LIS-catheters were used. **A)** Intensities of the 95% most abundant proteins were summed in a total proteome approach and compared between the UM-catheter and the LIS-catheter groups. **B)** A volcano plot for a subset of proteins. Negative mean rank difference indicates less protein on the LIS-catheter then on the UM-catheter and a significant difference is a -log10(P-value) over 1.3. The Fg chains (α-, β-, and γ-) are highlighted in green, serum albumin in orange, UDP-glucose 6-dehydrogenase, filamin-B and proteasome subunit beta type-5 in yellow.

## Discussion

This is the first study to show a diverse set of uropathogens including gram-negative, grampositive, and fungal species interact with Fg to more effectively bind to silicone urinary catheter surfaces. Furthermore, we found that by disrupting Fg deposition with LIS-catheters we reduced the ability of uropathogens to bind and colonize the catheter surface and bladder in an *in vivo* CAUTI mouse model. Moreover, LIS also reduced dissemination of *E. coli, P. aeruginosa and A. baumannii* into the kidneys and other organs. Finally, LIS-catheters didn’t increase inflammation and for half of the pathogens inflammation was reduced. Furthermore, the deposition of other host secreted proteins on LIS-catheters is around 6.5 fold less then UM-catheters. Together, these findings indicate that catheters made using LIS are a promising new antibiotic sparring approach for reducing or even preventing CAUTIs by interfering with protein deposition.

Pathogen-Fg interaction has been shown to be important during urinary catheterization for both *E. faecalis* and *S. aureus*. Binding to Fg is critical for efficient bladder colonization and biofilm formation on the catheter via protein-protein interaction using EbpA and ClfB adhesins, respectively and their disruption hinders colonization (A. L. Flores-Mireles et al., 2014; Walker et al., 2017). Gram-negative pathogens, *A. baumannii* and *P. mirabilis*, have shown colocalization with Fg during urinary catheterization (Gaston et al., 2020); however, the bacterial factors and any mode of interaction have not been described. While interaction of *E. coli* and *K. pneumoniae* with Fg during CAUTI has not been described, pathogenesis during urinary tract infection (UTI) (no catheterization) has been extensively studied. These studies show that type 1 pili, a chaperon-usher pathway (CUP) pili, allows them to colonize the bladder urothelium by binding to mannosylated receptors on the urothelial surface through the tip adhesin FimH (Klein & Hultgren, 2020). Furthermore, other CUP pili including the P pili, important for pyelonephritis, and the Fml pilus, important for colonizing inflamed bladder urothelium, bind specifically to sugar residues Galα1-4Gal in glycolipids and Gal(β1-3)GalNAc in glycoproteins, respectively (Klein & Hultgren, 2020). Interestedly, Fg is highly glycosylated, containing a wide variety of sugar residues including mannose, *N*-acetyl glucosamine, fucose, galactose, and N-acetylneuraminic acid (Adamczyk, Struwe, Ercan, Nigrovic, & Rudd, 2013). Therefore, the glycosylation in Fg may be recognized by CUP pili for colonization. Furthermore, *A. baumannii* CUP1 and CUP2 pili are essential for CAUTI, this together with its interactions with Fg *in vivo*, suggests that these pili may play a role in Fg interaction (Di Venanzio et al., 2019). Similarly, *P. aeruginosa* also encodes CUP pili, CupA, CupB, CupC and CupD, which have shown to be important for biofilm formation (Mikkelsen, Hui, Barraud, & Filloux, 2013); however, their contribution during CAUTI has yet to be investigated. Furthermore, *C. albicans* has several adhesins, ALS1, ALS3, and ALS9, which have a conserved peptide binding cavity shown to bind to Fg γ-chain (Hoyer & Cota, 2016).

Inhibition of initial uropathogen binding is crucial to reduce colonization and biofilm formation on urinary catheter surfaces and prevent subsequent CAUTI. To prevent surface binding, a variety of modified surfaces impregnated with antimicrobial or bacteriostatic compounds have been generated and, have proven to reduce microbial binding *in vitro* but not *in vivo* (Andersen & Flores-Mireles, 2019; Singha et al., 2017). It’s possible that *in vitro* studies do not efficiently mimic the complexities of the *in vivo* environment, for example; **1**) Growth media: the majority of the *in vitro* studies use laboratory rich or defined culture media, and it has been shown that laboratory media do not recapitulate the catheterized bladder environment that pathogens encounter (Colomer-Winter, Lemos, & Flores-Mireles, 2019; Xu et al., 2017). Specifically, urine culture conditions have shown to activate different bacterial transcriptional profiles than when grown in defined media (Conover et al., 2016; Xu et al., 2017), which may affect microbial persistence and survival. **2**) Host factors: host-secreted proteins are released into the bladder due to catheter-induced physical damage and subsequent inflammation. These proteins are deposited on the catheter surface and may hinder the release of antimicrobials or block interaction of antimicrobials with the pathogen. As has been observed with Fg deposition, host-protein deposition is not uniform, which may lead to antimicrobial release or pathogen-antimicrobial agent interaction at a sub-inhibitory concentrations (Di Venanzio et al., 2019; A. L. Flores-Mireles et al., 2014; A. L. Flores-Mireles et al., 2016; Walker et al., 2017). Consequently, these interactions can contribute to the development of multidrug-resistance among uropathogens (“Antibiotic resistance threats in the United States, 2019,” 2019).

Based on the role of deposited host-proteins in promoting microbial colonization, antifouling catheter coatings present a better approach to decreasing CAUTI prevalence rather than using biocidal or biostatic compounds, such as antibiotics, that promote resistance (Campoccia et al., 2013; Singha et al., 2017). Antifouling coatings are made from polymers and have shown resistance to protein deposition (He et al., 2016); however, these coatings can become unstable over time and be difficult to produce (Singha et al., 2017). Most reports on the use of purely antifouling coatings to combat CAUTI have shown a successful reduction in bacterial colonization *in vitro* yet have not been tested *in vivo* (Andersen & Flores-Mireles, 2019). This may be partly explained by the fact that many of the antifouling coatings are optimized to target bacterial adhesion, as it is understood that the first stage of biofilm development is bacterial attachment to a surface (Faustino et al., 2020). However, in a complex environment such as the *in vivo* bladder, the first change to the catheter surface is the adhesion of a complex set of host-generated proteins and biological molecules, generally referred to as a conditioning film, which can mask the surface (Faustino et al., 2020; Gaston et al., 2020; Scotland, Kung, Chew, & Lange, 2020). Yet studies on catheter coatings to-date have rarely focused on the role of the host in infection establishment; namely, the host-secreted proteins. The data we present here suggest that this missing element may at least partly explain the differing results seen for most antifouling catheter treatments *in vitro* vs *in vivo*.

This study used clinically relevant silicone oil to create a simple liquid-infused polymer that was not only bacteria-resistant but also protein-resistant, filling in the missing link between *in vitro* and *in vivo* work, the conditioning film. Furthermore, lack of an exacerbated inflammation response seen in our results echoes previous *in vivo* work showing reduced capsule formation in implants coated with a liquid layer, suggesting that the use of a liquid surface may convey additional antiinflammatory benefits (Chen et al., 2017).

A deeper understanding of the pathogenesis of CAUTI is critical to moving beyond current developmental roadblocks and create more efficient intervention strategies. Here, we have shown that that infusion of silicone with an immiscible liquid coating significantly decreases Fg deposition and microbial binding by using *in vitro* conditions that more thoroughly recapitulate the catheterized bladder environment. Importantly, our *in vitro* results were confirmed *in vivo* using our established mouse model of CAUTI. Our data showed that LIS-catheters are refractory to bacterial colonization without targeting microbial survival, which often leads to antimicrobial resistance, and thus holds tremendous potential for the development of lasting and effective CAUTI treatments. These types of technologies are desperately needed to achieve better public health by decreasing healthcare-associated infections and promoting long-term wellness.

## Data and materials availability

The data that support the findings of this study are either provided in the source data or are available from the corresponding authors upon reasonable request. RAW and processed MS-MS/MS data are available for review at http://mchampion-nas.esc.nd.edu:5000/sharing/oOBGtDP6I Password: CathRAW21*. Additionally, these data will be transferred to the MassIVE (https://massive.ucsd.edu/ProteoSAFe/static/massive.jsp) public repository after review and this link will be updated with an accession number. This study did not generate new unique reagents.

## Acknowledgments

We thank members of the Flores-Mireles and Howell laboratories for their helpful suggestions and making this project possible. Special thank you to Dr. Matthew Champion for his insight into proteomic analysis and the ND CORE Facilities for their work on the Mass Spectrometer and tissue processing. Thank you to Mario Feldman and Gisela Di Venanzio for providing the *A. baumannii* strain and antibodies. This work was funded by the University of Notre Dame Institutional Funds and seed grant FY19SEED6 (to A.L.F.M., A.A.L.B. and A.M.), National Institute of Health grant R01-DK128805 (to A.L.F.M., M.J.A., C.H. and C.K.F.) and National Science Foundation grant no CBET-2029378 (to C.H. and C.K.F.).

## Author Contributions

Conceptualization: MJA, CKF, CH, ALFM. Methodology: MJA, CKF, CH, ALFM, MMC. Investigation: MJA, CKF, AALB, AM, MMC. Visualization: MJA, ALFM. Funding acquisition: ALFM, CH. Project administration: ALFM, CH. Supervision: ALFM, CH. Writing – original draft: MJA. Writing – review & editing: MJA, ALFM, CKF, CH, MMC

## Declaration of interests

The authors declare no competing financial interests.

## Materials and Methods

### Mouse Infection Models

Mice used in this study were ~6-week-old female wild-type C57BL/6 mice purchased from Jackson Laboratory and The National Institute of Cancer Research. Mice were subjected to transurethral implantation and inoculated as previously described.(Conover, Flores-Mireles, Hibbing, Dodson, & Hultgren, 2015) Briefly, mice were anesthetized by inhalation of isoflurane and implanted with a 6-mm-long UM-silicone or LIS-catheter. Mice were infected immediately following catheter implantation with 50 μl of ~2 × 10^7^ CFU/mL in PBS introduced into the bladder lumen by transurethral inoculation (unless otherwise noted (ST1). For all mouse experiments microbes were grown in their corresponding media (ST1). To harvest the catheters and organs, mice were sacrificed at 24hrs post infection by cervical dislocation after anesthesia inhalation; the silicone catheter, bladder, kidneys, heart and spleen were aseptically harvested. Catheters were either subjected to sonication (Branson, Ultrasonic Bath) for CFU enumeration analysis, fixed for imaging via standard IF procedure described above, or sent for proteomic analysis as described below using nonimplanted catheters as controls. Bladders for immunofluorescence and histology were fixed and processed as described below. Kidneys, Spleens and Hearts were all used for CFU analysis. The University of Notre Dame Institutional Animal Care and Use Committee approved all mouse infections and procedures as part of protocol number 18-08-4792MD. All animal care was consistent with the Guide for the Care and Use of Laboratory Animals from the National Research Council.

### Bladder IHC and H&E staining of mouse bladders

Mouse bladders were fixed in formalin overnight, before being processed for sectioning and staining as previously described (Walker et al., 2017). Briefly, bladder sections were deparaffinized, rehydrated, and rinsed with water. Antigen retrieval was accomplished by boiling the samples in Na-citrate, washing in water, and then incubating in PBS three times. Sections were then blocked (1x PBS, 1.5% BSA, 0.1% Sodium Azide), washed in PBS, and incubated with appropriate primary antibodies overnight at 4 °C. Next, sections were washed with PBS, incubated with secondary antibodies for 2 h at RT, and washed once more in PBS prior to Hoechst dye staining. Hematoxylin and Eosin (H&E) stain for light microscopy was done by the CORE facilities at the University of Notre Dame (ND CORE). All imaging was done using a Zeiss inverted light microscope (Carl Zeiss, Axio Observer). Zen Pro (Carl Zeiss, Thornwood, NY) and ImageJ software were used to analyze the images.

### Human Urine Collection

Human urine was collected and pooled from at least two healthy female donors between 20–40 years of age. Donors had no history of kidney disease, diabetes or recent antibiotic treatment. Urine was sterilized using a 0.22μm filter (Sigma Aldrich) and pH 6.0–6.5. When supplemented with Bovine Serum Albumin (BSA) (VWR Lifesciences), urine was filter sterilized again following BSA addition. All participants signed an informed consent form and protocols were approved by the local Internal Review Board at the University of Notre Dame under study #19-04-5273.

### Microbial Growth Conditions in Supplemented Urine

*E. faecalis*, and *C. albicans* were grown static for ~5 hrs in 5 mL of respective media **(Table S2)** followed by static overnight culture in human urine supplemented with 20mg/mL BSA (urine BSA20). *E. coli, K. pneumoniae, P. mirabilis, A. baumanii* and *P. aeruginosa* were grown 5hrs shaking at 37 °C in LB then static for 24 hours, supplemented into fresh urine BSA for another 24 hours static (2×24hrs) in urine BSA20. All cultures were washed in PBS (Sigma) 3 times and resuspended in assay appropriate media.

### Silicone Disk Preparation

8mm disks of UM-silicone (Nalgene 50 silicone tubing, Brand Products) or LIS were cut using a leather hole punch. UM disks were washed 3 times in PBS and air dried. LIS disks were stored in filter sterilized modifying liquid at RT. Disks were skewered onto needles (BD) to hold them in place and put in 5mL glass tubes (Thermo Scientific) or placed on the bottom of 96 well plate wells (Fisher Scientific) (UM silicone only). Plates and glass tubes were UV sterilized for >30min prior to use.

### Protein Binding Assays

Human fibrinogen free from plasminogen and von Willebrand factor (Enzyme Research Laboratory #FB3) was diluted to 150ug/mL in PBS. 500uL of 150ug/mL Fg was added to each disk in glass tubes, sealed, and left over night at 4 °C. Disks were then processed according to standard immunofluorescence (IF) procedure as previously described.(Colomer-Winter et al., 2019) Briefly, disks were washed 3 times in PBS, fixed with 10% Neutralized formalin (Leica), blocked, and stained using Goat anti-Fg primary antibody (Sigma) (1:1000) and Donkey anti-Goat IRD800 secondary antibody (Invitrogen) (1:1000). Disks were then dried over night at 4°C and imaged on an Odyssey Imaging System (LI-COR Biosciences) to examine the infrared signal. Intensities for each catheter piece were normalized against a negative control and then made relative to the pieces coated with Fg which was assigned to 100%. Images were processed using Image Studio Software (LI-COR, Lincoln, NE) Microsoft Excel and graphed on GraphPad Prism (GraphPad Software, San Diego, CA).

### Microbial Binding Assays

For assessing the effect of protein deposition on microbial binding, 100uL of 150ug/mL Human Fg, 100uL of 150ug/mL BSA, or 100uL of PBS were incubated on UM-silicone disks in 96 well plates overnight at 4°C. The following day disks were washed 3 times with PBS followed by a 2hr RT incubation in 100uL of urine containing microbes at a concentration of ~10^8^ CFU/mL.

For assessing microbial binding to UM-silicone versus LIS, 500uL of microbe containing media was added to prepared disks in glass tubes. Standard IF procedure was then followed as described above using goat anti-Fg and rabbit anti-microbe primary antibodies (1:1000) (see ST1 for details). Secondary antibodies used were Donkey anti-Goat IRD800 and Donkey anti-Rabbit IRD680 (1:5000). Quantification of binding was done using ImageStudio Software (LI-COR). Intensities for each catheter piece were normalized against a negative control and then made relative to the pieces coated with Fg which was assigned to 100%.

### Weight measurement of silicone tubes and Tygon tubes versus infusion time

Five samples of 20 cm Tygon tube (14-171-219, Saint-Gobain Tygon S3 TM 3603 Flexible Tubings, Fisher Scientific, USA) or silicone tube (8060-0030, NalgeneTM 50 Platinum-cured Silicone Tubing, Thermo Scientific, USA) were utilized in weight measurement. Weight of the tubes prior to infusion were measured with an analytical balance (AL204, Analytical Balance, Mettler Toledo, Germany), results were marked as “0 h infused in silicone oil”. After the measurement of the initial weights, the tubes were incubated with silicone oil (DMS-T15, Polydimethylsiloxane, trimethylsiloxy, 50 cSt, GelestSInc., USA) until designated time points. For each time point, tubes were removed from the oil with forceps and held vertically for 30 seconds for the excess silicone oil to flow out of the tube. The bottoms of the tubes were then gently dabbed with Kimwipes™ (Kimwipe, Kimberly-Clark Corp., USA), and then subjected to weight measurement. After measurement, the tubes were placed back into silicone oil until the next time point. Tubes were measured every 3 hrs for the first 2 days; every 6 hrs from day 3 to day 6; and every 24 hrs from day 6 and onwards. Measurements were taken until data showed no significant increase, and that the plateau trendline consist of at least 3 data points.

### Weight measurement of mouse catheters versus infusion time

Five samples of 20 cm mouse catheter (SIL 025, RenaSil Silicone Rubber Tubing, Braintree Scientific, Inc., USA) were utilized in weight measurement. Weight of the tubes prior to infusion were measured with an analytical balance, results were marked as “0 min infused in silicone oil”. After the measurement of the initial weights, the tubes were incubated with silicone oil until designated time points. For each time point, catheters were removed from the oil with forceps, a Kimwipes™ was immediately pressed against the bottom of the catheters to remove the excess silicone oil via capillary action. After the excess oil was drained, catheters were then subjected to weight measurement. The catheters were placed back into silicone oil until the next time point. Catheters were measured every 1 minute for the first 5 minutes of the experiment; every 2 minutes from 5-15 minutes; and every 5 minutes from 15 minutes and onwards. Measurements were taken until data showed no significant increase, and that the plateau trendline consist of at least 3 data points.

### Parameter measurement of silicone tube before and after infusion

The length, inner diameter and outer diameter of the silicone tubes were measured before silicone oil infusion and after complete infusion (after incubating with silicone oil for >7 days). All parameters were measured using a digital caliper (06-664-16, Fisherbrand™ Traceable™ Digital Calipers, Fisher Scientific, USA).

### Parameter measurement of mouse catheters before and after infusion

The length, inner diameter, and outer diameter of the mouse catheters were measured before silicone oil infusion and after complete infusion (after incubating with silicone oil for >30 minutes). The length of the mouse catheter was measured using a digital caliper. Photos of the tube openings of the catheters and a scale of known length were taken. The inner and outer diameter were then estimated via ImageJ. Percentage weight change of Tygon tube, silicone tube and mouse catheters were calculated based on the formula below:

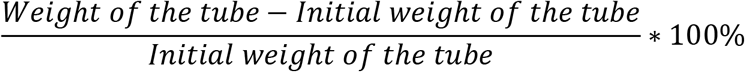

### Proteomic analysis of mouse catheters

Five mice were catheterized with a LIS-catheter, 4 mice were catheterized with an UM-catheter and catheters harvested 24hrs after infection with *E. faecalis* OG1RF. Harvested catheters were put into 100uL of SDS buffer (100mM Tris HCl pH-8.8, 10mM DTT and 2% SDS), then vortexed for 30 sec, heated for 5 min at 90 °C, sonicated for 30min and the process repeated once more. Samples were sent to the Mass Spectrometry and Proteomics Facility at Notre Dame (MSPF) for proteomic analysis. Proteins were further reduced in DTT, alkylated and digested with trypsin using Suspension Trap and protocols.(Zougman, Selby, & Banks, 2014) nLC-MS-MS/MS was performed essentially as described in (Sanchez et al., 2020) on a Q-Exactive instrument (Thermo). Proteins were identified and quantified using MaxLFQ (Label Free Quantification) within MaxQuant and cutoff at a 1% FDR.(Cox & Mann, 2008) This generated a total of 8 data records from UM-catheters and 10 from LIS-catheters. Data reduction was performed by removing contaminants proteins. Protein abundance for each catheter type was then calculated by summing the LFQ intensity of proteins which comprised 95% of the total abundance on the catheters. Strict filtering criteria of at least 2 replicates with technical replication from the UM-catheters and 3replicates with technical duplication from the LIS-catheters were required to keep an identification. Abundance of the reduced proteins was plotted using Graph Pad Prism. Statistical significance was tested using Mann-Whitney *U*. A volcano plot was created using the ranked mean difference for each protein and -log of calculated P-values with an alpha =0.05.

### Statistical Analysis

Unless otherwise stated, data from at least 3 experiments were pooled for each assay. Significance of experimental results were assessed by Mann-Whitney *U* test using GraphPad Prism, version 7.03 (GraphPad Software, San Diego, CA). Significance values on graphs are *p≤ .05, **p≤ .01, ***p≤ .001 and ****p≤ .00001.

### Antibodies used in this study

Primary antibodies against microbial pathogens used in the study are listed in table S2. Primary antibodies against non-pathogens; Fg, Goat anti-Fibrinogen and neutrophils, Rat anti-Ly6G. Secondary antibodies used for IF in the study: IRDye 800CW donkey anti goat (LI-COR) and IRDye 680LT Donkey anti-rabbit (LI-COR). Secondary antibodies for IHC; Donkey anti-goat (Life Technologies Corporation), Donkey anti-rabbit (Invitrogen), Donkey anti-mouse (Invitrogen) and Donkey anti-rat (Invitrogen).

## Supplementary Figures

**Fig. S1.**
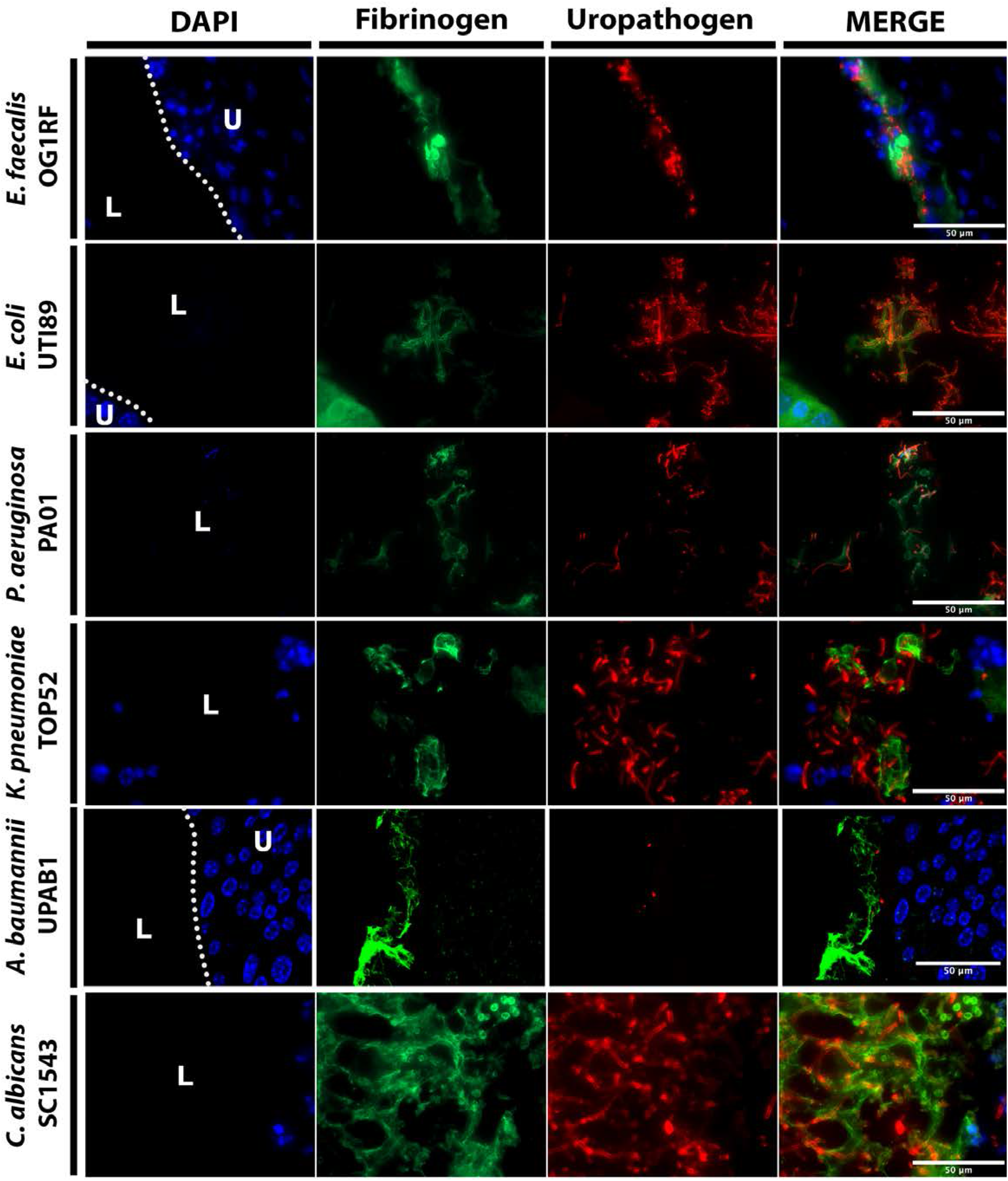
Montages of Figure 1 merged images. Mice were implanted and infected with 1× 10^6^ CFU of the respective uropathogens. At 24 hpi, bladders tissues were harvested, fixed, and parafilm-embedded. Bladder were subjected to IF analysis, antibody staining was used to detect Fg (anti-Fg; green), uropathogens (red), and DAPI (blue) for cell nuclei. Scale bars, 50 um. Magnification 100x.

**Fig. S2.**
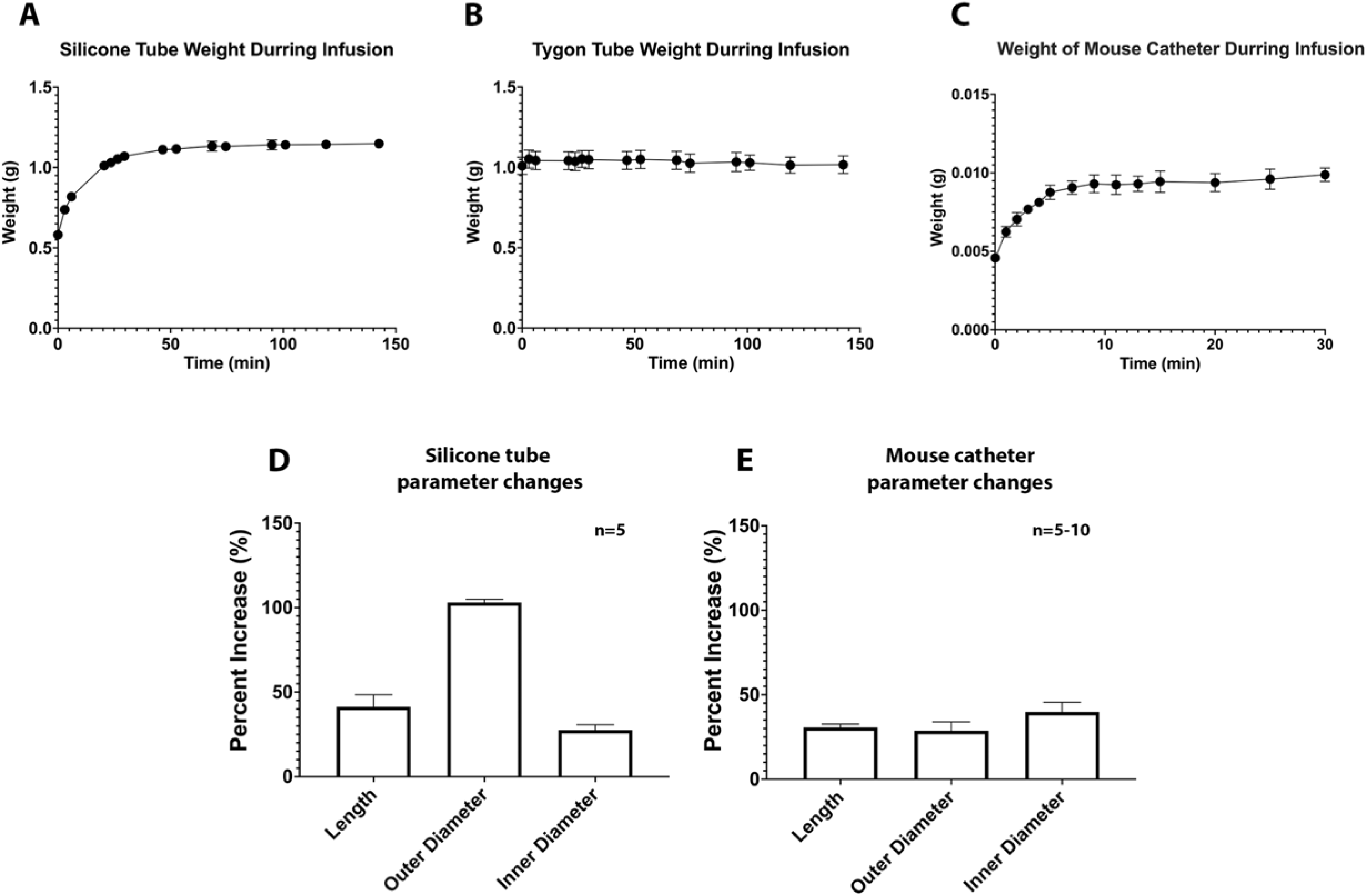
Characterization of LI tubing and mouse catheters. **A)** Weight of silicone tubes were measured in designated time points before and during silicone oil infusion, the mean (±SEM) of n = 5 silicone tubes over infusion time was shown in this figure. **B)** Weight of Tygon tubes were measured in designated time points before and during silicone oil infusion, the mean (±SEM) of n = 5 silicone tubes over infusion time was shown in this figure. **C)** Weight of mouse catheters were measured in designated time points before and during silicone oil infusion, the mean (±SEM) of n = 5 mouse catheter over infusion time was shown in this figure. The length, outer and inner diameter of **F)** silicone catheters (n = 5) or **G)** mouse catheters (n = 5-10) were measured before and after infusion and the percentage change was calculated.

**Figure S3.**
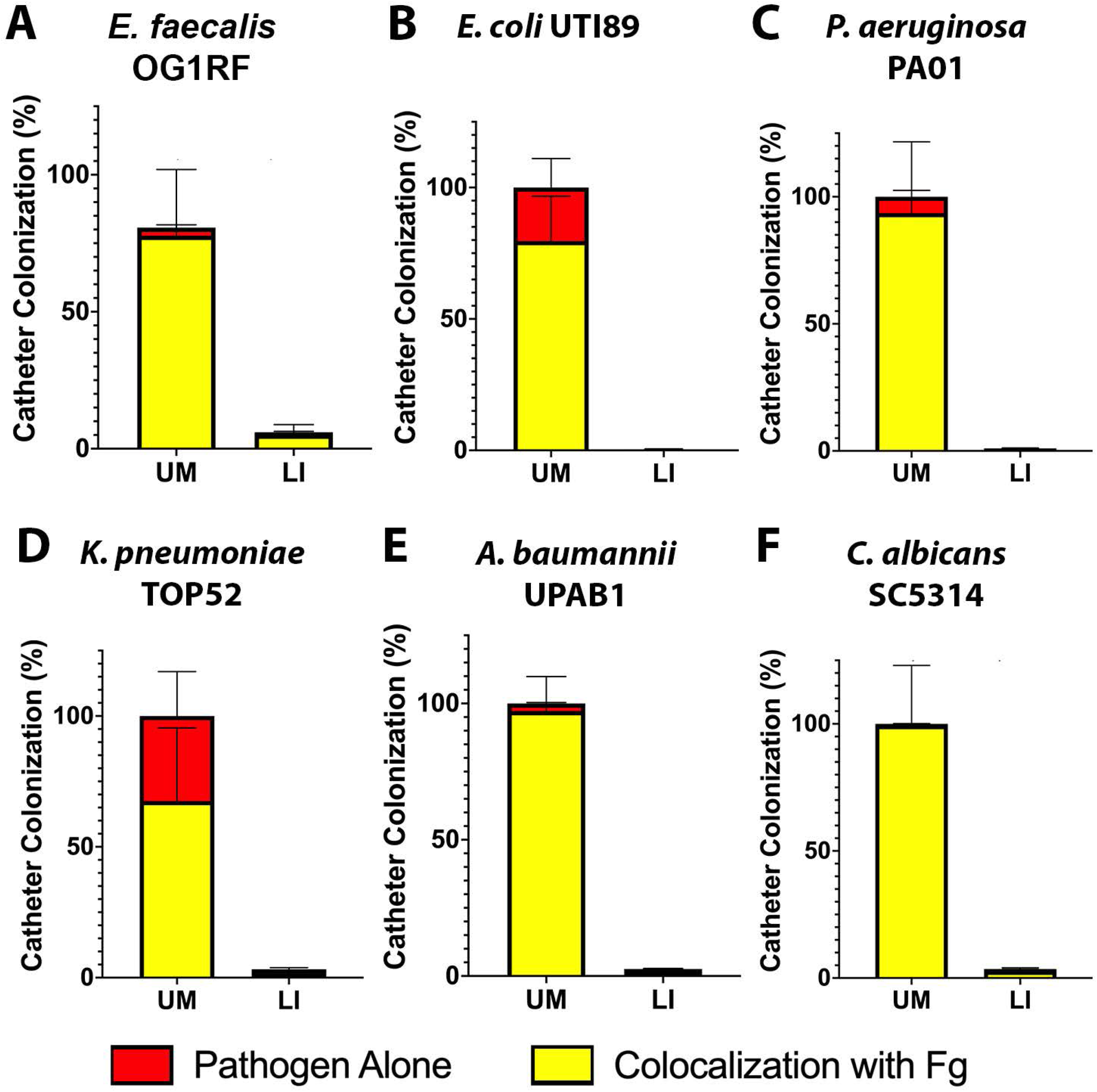
Pathogen predilection for fibrinogen. Quantification of uropathogen-Fg colocalization on UM and LIS-catheters from mice catheterized and infected with one of six uropathogens. Quantification was done using pixel color counter from Fiji where colocalization (yellow) of Fg (green) and pathogen (red) were quantified and compared to the total pathogen colonization of the catheter.

**Fig. S4.**
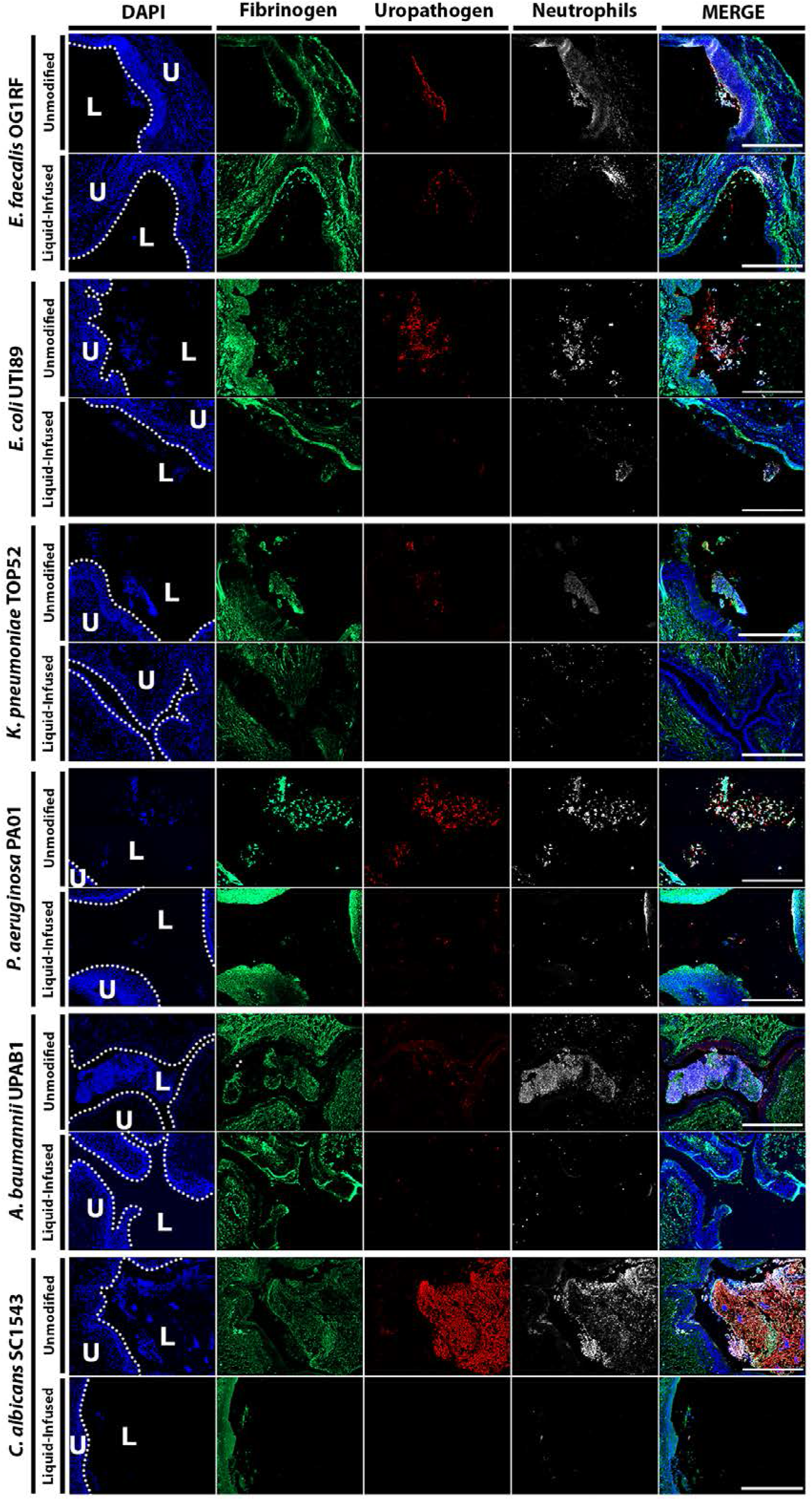
Montage of Figure 5 merged images. Mice were implanted with either an unmodified catheter or a liquid-infused catheter and infected with 1× 10^6^ CFU of the respective uropathogens. At 24 hpi, bladders tissues were harvested, fixed, and parafilm-embedded. Bladder were subjected to IF analysis, antibody staining was used to detect Fg (anti-Fg; green), uropathogens (red), neutrophils (anti-Ly6G; white) and DAPI (blue) for cell nuclei. Scale bars, 500 um. Images are stitched 2×2 tiles at 20x magnification.

**Table S1. List of proteins found on LI and UM mouse catheters infected with *E. faecalis* OG1RF.** The average number of peptides for each protein found on 10 mouse catheters sorted by greatest abundance on the UM catheter.

**Table S2. Microbe details.** List of microbial strains and their corresponding growth conditions, inoculum concentrations and antibodies used in this study

## Notes

### Competing Interest Statement

The authors have declared no competing interest.

